# Outpatient antibiotic prescribing and demographic factors associated with state-level septicemia mortality rates in US adults

**DOI:** 10.1101/480137

**Authors:** Edward Goldstein, Marc Lipsitch

**Affiliations:** Center for Communicable Disease Dynamics, Department of Epidemiology, Harvard TH Chan School of Public Health, Boston, MA 02115 USA; Department of Immunology and Infectious Diseases, Harvard TH Chan School of Public Health, Boston, MA 02115 USA

## Abstract

**Background:** Rates of septicemia/sepsis mortality and hospitalization in the US have risen significantly during the recent years, and antibiotic use may contribute to those rates through various mechanisms.

**Methods:** We used multivariable linear regression to relate state-specific rates of outpatient prescribing overall for fluoroquinolones, penicillins, macrolides, and cephalosporins between 2013-2014 to state-specific rates of septicemia mortality (ICD-10 codes A40-41 present as either the underlying or contributing causes of death) in each of the following age groups of adults: (18-49y, 50-64y, 65-74y, 75-84y, 85+y) between 2013-2014, adjusting for median household income, average annual temperature, age-specific percentages of state residents who (i) lived below the poverty level, (ii) were African American, (iii) lacked health insurance (in non-elderly age groups), and random effects associated with the different US Health and Human Services (HHS) regions.

**Results:** Rates of penicillin prescribing were positively associated with septicemia mortality rates in the analyses for persons aged 65-74y, 75-84y and over 85y. Rates of cephalosporin prescribing were positively associated with septicemia mortality rates in the analyses for persons aged 18-49y and 65-74y. Rates of fluoroquinolone prescribing were positively associated with septicemia mortality rates in the analyses for persons aged 18-49y. Percent African Americans in a given age group was positively associated with septicemia mortality rates in the analyses for age groups over 65y, and 18-49y. Percent of residents in a given age group living below the poverty level was positively associated with septicemia mortality rates in the analysis for persons aged 65-74y.

**Conclusions:** Our results suggest that rates of penicillin prescribing are associated with rates of septicemia mortality in older US adults, while rates of cephalosporin prescribing are associated with rates of septicemia mortality in persons aged 18-49y and 65-74y, and rates of fluoroquinolone prescribing are associated with rates of septicemia mortality in persons aged 18-49y. Further studies are needed to better understand the potential effect of antibiotic replacement in the treatment of different syndromes, such as replacement of fluoroquinolones by other antibiotics, possibly penicillins and cephalosporins following the recent US FDA guidelines on restriction of fluoroquinolone use, on the rates of sepsis mortality.

## Introduction

Rates of hospitalization with septicemia and sepsis, associated mortality and monetary costs have been rising rapidly during the past decades in the US [1-4]. Moreover, the US CDC estimate of 270,000 annual deaths in the US resulting from sepsis [5] is expected to increase significantly if longer-term (e.g. 90-day) mortality following hospitalization with septicemia/sepsis were accounted for [6]. While changes in diagnostic practices contributed to the growth in the volume of hospitalizations with septicemia/sepsis in the diagnosis [7,8], those changes cannot fully explain the rise in the rates of hospitalization with septicemia/sepsis, particularly prior to 2010 [9]. Indeed, trends in the rates of US hospitalizations with any diagnosis of sepsis between 2003-2009 closely resemble the trends in the rates of hospitalizations that involved infection and the use of mechanical ventilation (Figure 1 in [9]). Use of antibiotics and antibiotic resistance may also contribute to the rates of hospitalization with septicemia and the associated mortality [10-13]. Antibiotic resistance and use can facilitate the progression to a severe disease state when infections not cleared by antibiotics prescribed during outpatient and/or the inpatient treatment may eventually devolve into sepsis, e.g. [14,15]. Antibiotic use may also contribute to prevalence of infections resistant to other antibiotics that may lead to severe outcomes. For example, fluoroquinolone use was found to be associated with methicillin-resistant *S. aureus* (MRSA) infections [16,17], while amoxicillin use was found to be associated with trimethoprim resistance in Enterobacteriaceae [18], with trimethoprim-resistant urinary tract infections (UTIs) leading to bacteremia outcomes [15].

Our previous work had suggested associations between the use of penicillins and fluoroquinolones, antibiotic resistance, and rates of septicemia hospitalization in US adults [10,11,19]. In this paper we study the relation between the rates of outpatient prescribing for different antibiotic classes and the rates of mortality with septicemia (ICD-10 codes A40-41 present as either the underlying or contributing causes of death) in US adults. We note that the relation between outpatient antibiotic prescribing and the rates of septicemia hospitalization may be different from the relation between outpatient antibiotic prescribing and the rates of septicemia mortality. Indeed, case-fatality rates for hospitalizations with septicemia depend on the types of infection leading to septicemia, including their antibiotic resistance profiles [12,13], demographic factors, including presence of comorbidities [13,20], practices for antibiotic prescribing in the treatment of inpatients with septicemia [12], etc. Moreover, major differences exit in coding practices for a septicemia diagnosis in hospitalized patients, with those differences further contributing to differences in case fatality rates for inpatients with septicemia in the hospitalization diagnosis. For example, a study from Kaiser Permanente Northern California suggested that in-hospital fatality rates for their inpatients with hospitalization codes for septicemia/sepsis were significantly lower compared to the NIS data [21]. In this study, we use a multivariable regression framework to relate the rates of outpatient prescribing of fluoroquinolones, penicillins, cephalosporins and macrolides to the rates of septicemia mortality in different age groups of US adults, adjusting for additional effects. We hope that such ecological analyses can lead to further study of the contribution of different antibiotics to the rates of sepsis mortality, including the utility of a shift in prescribing from some antibiotics to certain others in the treatment of different syndromes for reducing the levels of sepsis mortality in the US.

## Methods

### Data

We extracted data on annual state-specific mortality with septicemia (ICD-10 codes A40-A41 representing either the underlying or a contributing cause of death) between 2013-2014 for different age groups of adults (18-49y, 50-64y, 65-74y, 75-84y, 85+y) from the US CDC Wonder database [22]. We note that for the majority of those deaths, septicemia is listed as a contributing, rather than the underlying cause of death on the death certificate [23]. Data on the annual state-specific per capita rates of outpatient prescribing for four classes of antibiotics: fluoroquinolones, penicillins, macrolides, and cephalosporins in 2013 and 2014 were obtain from the US CDC Antibiotic Patient Safety Atlas database [24]. Data on median household income for US states between 2013-2013 were extracted from [25]. Data on average monthly temperature for US states were obtained from [26]. Annual state-specific population estimates, including ones of African American populations in different age groups were extracted from [27]. Data on the percent of state residents who lived below the poverty level, and those who lacked health insurance were extracted form the US Census Bureau database [28].

### Regression model

For each age group of adults, (18-49y, 50-64y, 65-74y, 75-84y, 85+y), we applied multivariable linear regression to relate the average annual state-specific outpatient prescribing rates (per 1,000 state residents) for fluoroquinolones, penicillins, macrolides, and cephalosporins between 2013-2014 to the average annual state-specific rates of septicemia mortality per 100,000 individuals in a given age group between 2013-2014 (dependent variable). Besides the antibiotic prescribing rates, the other covariates were the state-specific median household income, percentages of state residents in a given age group who lived below the poverty level, those who were African American, those who lacked health insurance (in the non-elderly age groups, as health insurance, particularly Medicare coverage levels in the elderly are very high), as well as the state-specific average annual temperature. We note that septicemia mortality rates in African Americans are elevated [29]. We also note that temperature may influence bacterial growth rates and/or transmission mediated effects [30], which in turn may affect both the prevalence of antibiotic resistance [30,19], and the acquisition/severity of bacterial infections. To adjust for additional factors not accounted for by the covariates used in the model, we include random effects for the ten Health and Human Services (HHS) regions in the US. Specifically, for each state *s*, let *MR*(*s*) be the average annual state-specific rate of mortality (per 100,000) with septicemia in the given age group between 2013-2014, *A_i_*(*s*) (*i=*1,..,4) be the average annual state-specific outpatient prescribing rates, per 1,000 state residents, for the four studied classes of antibiotics between 2013-2014; *I*(*s*) be the median state-specific household income between 2013-2014; *T*(*s*) be the state-specific average annual temperature (°F) between 2005-2014; *AA*(*s*) be the age-specific percent of state residents between 2013-2014 who were African American; *BPL*(*s*) be the age-specific percent of state residents between 2013-2014 who lived below the poverty level; *LHI*(*s*)be the age-specific percent of state residents who lacked health insurance between 2013-2014 (for non-elderly age groups); and *RE*(*s*) be the random effect for the corresponding HHS region. Then

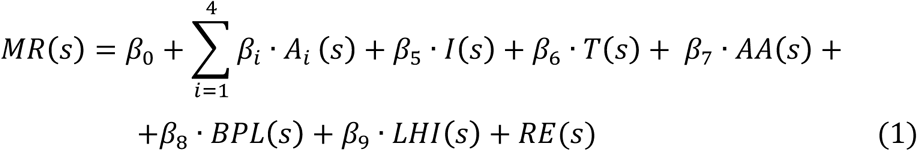

## Results

Table 1 shows the septicemia mortality rates (per 100,000) in the five age groups of adults (18-49y, 50-64y, 65-74y, 75-84y, 85+y) between 2013-2014 for the 50 US states and the District of Columbia (median among state-specific rates + lower/upper quartiles). Those rates grow rapidly with age, with the rates in younger adults being by far the lowest (median=7.3), and highest in persons aged over 85 years (median = 745.8). Table 1 also shows, for each of the five age groups of adults, the rates of septicemia mortality between 2013-2014 in each of the ten US Health and Human Services (HHS) regions. The septicemia mortality rates in all age groups over 50 years are highest in HHS region 6 (South Central); for age groups over 75 years, the second highest rates are in HHS region 2 (NY + NJ); for age groups 65-74 years and 50-64 years, the second highest rates are in HHS region 3 (Mid Atlantic, including VA + WV) and 4 (Southeast) correspondingly. The lowest mortality rates in all age groups are in HHS region 8 (Mountain), followed by region 10 (Northwest) for persons aged over 85 years, region 7 (IA, KS, MO, NE) for persons aged 75-84 years and region 10 (Northwest) for persons aged 65-74 years.

**Table 1:** Septicemia mortality rates (per 100,000) in the five age groups of adults (18-49y, 50-64y, 65-74y, 75-84y, 85+y) between 2013-2014 for (i) the 50 US states and the District of Columbia (median among state-specific rates + lower/upper quartiles); (ii) the 10 US HHS regions.

|  | 18-49y | 50-64y | 65-74y | 75-85y | 85+y |
| --- | --- | --- | --- | --- | --- |
| 50 states +DC | 7.3<br>[6.1,9.3] | 51.3<br>[39.7,64.5] | 136<br>[109.8,166.4] | 339.9<br>[253.3,381.4] | 745.8<br>[580,862.4] |
| HHS Region 1 | 6.1 | 39.9 | 128.7 | 326.6 | 763.8 |
| HHS Region 2 | 6 | 47.4 | 144.2 | 382.5 | 919.8 |
| HHS Region 3 | 7.7 | 56.6 | 156.3 | 373.4 | 828 |
| HHS Region 4 | 10 | 63.7 | 150.9 | 336.8 | 736.2 |
| HHS Region 5 | 7.1 | 49.4 | 136.6 | 332.3 | 697.4 |
| HHS Region 6 | 9.8 | 71.2 | 180.6 | 410.5 | 946.8 |
| HHS Region 7 | 6.1 | 48 | 122.3 | 277.6 | 630.5 |
| HHS Region 8 | 5.7 | 36.9 | 100 | 251.2 | 566.4 |
| HHS Region 9 | 7.4 | 52.2 | 131.3 | 312.7 | 746.9 |
| HHS Region 10 | 6 | 43.8 | 109.2 | 284.4 | 642.2 |

Table 2 shows the estimates, for each age group of US adults, of the regression coefficients for the different covariates in the regression model (eq. 1). Those regression coefficients estimate the change in septicemia mortality rates (per 100,000 individuals in a given age group) when the corresponding covariate increases by 1. Table 2 suggests that rates of outpatient penicillin prescribing were positively associated with septicemia mortality rates in the analyses for persons aged 65-74 years, 75-84 years, and over 85 years. Rates of outpatient cephalosporin prescribing were positively associated with septicemia mortality rates in the analyses for persons aged 18-49 years and 65-74 years. Rates of outpatient fluoroquinolone prescribing were positively associated with septicemia mortality rates in the analyses for persons aged 18-49 years. Percent of African Americans among state residents in a given age group was positively associated with septicemia mortality rates in the analyses for persons aged 18-49 years, 65-74 years, 75-84 years, and over 85 years. Percent of state residents in a given age group living below the poverty level was positively associated with septicemia mortality rates in the analysis for persons aged 65-74y. Median household income was negatively associated with septicemia mortality rates in the analysis for persons aged 18-49 years.

**Table 2:** Estimates of the regression coefficients for the different covariates in the inference model (eq. 1). Each coefficient estimates the change in septicemia mortality rates (per 100,000 individuals in a given age group) when the corresponding covariate increases by 1.

|  | Aged<br>18-49y | Aged<br>50-64y | Aged<br>65-74y | Aged<br>75-84y | Aged<br>85+y |
| --- | --- | --- | --- | --- | --- |
| Fluoroquinolones<br>(prescription per<br>1000 residents/y) | **0.05**<br>**(0,0.1)** | 0.2<br>(-0.1,0.49) | -0.33<br>(-0.96,0.29) | -0.33<br>(-2.1,1.4) | -3.4<br>(-7.9,1.2) |
| Penicillins<br>(prescription per<br>1000 residents/y) | 0<br>(-0.03,0.02) | 0.13<br>(-0.01,0.28) | **0.4**<br>**(0.11,0.69)** | **1.2**<br>**(0.38,1.9)** | **3.4**<br>**(1.5,5.4)** |
| Cephalosporins<br>(prescription per<br>1000 residents/y) | **0.03**<br>**(0,0.06)** | 0.11<br>(-0.05,0.27) | **0.51**<br>**(0.15,0.87)** | 0.25<br>(-0.72,1.2) | 0.07<br>(-2.3,2.5) |
| Macrolides<br>(prescription per<br>1000 residents/y) | -0.02<br>(-0.05,0.01) | -0.09<br>(-0.26,0.07) | -0.13<br>(-0.48,0.23) | 0.17<br>(-0.8,1.1) | 1.2<br>(-1.2,3.7) |
| Median household<br>income (\$1000) | **-0.09**<br>**(-0.17,-0.02)** | -0.07<br>(-0.45,0.32) | -0.09<br>(-0.74,0.57) | 0.75<br>(-1.1,2.6) | 0.11<br>(-4.4,4.6) |
| Average annual<br>temperature (°F) | -0.07<br>(-0.14,0.01) | 0.06<br>(-0.27,0.39) | 0.26<br>(-0.46,0.98) | -0.48<br>(-2.4,1.4) | 1<br>(-3.9,6) |
| Percent African<br>Americans | **0.08**<br>**(0.02,0.15)** | 0.3<br>(-0.01,0.61) | **1.3**<br>**(0.54,2)** | **2.8**<br>**(1.1,4.5)** | **6.1**<br>**(1.9,10.4)** |
| Percent living<br>below poverty level | -0.09<br>(-0.26,0.09) | 0.32<br>(-0.71,1.4) | **2.1**<br>**(0.33,3.9)** | -1.5<br>(-4.8,1.9) | -7.1<br>(-15.4,1.2) |
| Percent lacking<br>health insurance | 0<br>(-0.1,0.1) | 0.33<br>(-0.37,1) | X | X | X |

## Discussion

Rates of septicemia/sepsis hospitalization and associated mortality in the US have risen during the recent years [1,2]. While outpatient antibiotic prescribing can contribute to progression to septicemia hospitalization (Introduction), and affect the resistance profiles for infections in hospitalized patients, which in turn affect their likelihood of survival [12,13], there is limited information on the strength of the relation between the levels of outpatient prescribing of different antibiotics and septicemia mortality rates in the US. In this study, we used a multivariable regression framework to relate the rates of outpatient prescribing of fluoroquinolones, penicillins, cephalosporins and macrolides to septicemia mortality rates in different groups of US adults. We have found that rates of penicillin prescribing are associated with septicemia mortality rates in older adults, which agrees with our earlier finding about the association between the rates of penicillin prescribing and the rates of septicemia hospitalization in older US adults [10]. We also note the high prevalence of resistance to penicillins in both the Gram-negative and Gram-positive bacteria (e.g. [31-33]). We have also found an association between the rates of cephalosporin prescribing and the rates of septicemia mortality in persons aged 18-49 years and 65-75 years, as well as the rates of fluoroquinolone prescribing and the rates of septicemia mortality in persons aged 18-49 years. We note that prevalence of cephalosporin resistance and the frequency of extended-spectrum beta-lactamase (ESBL) production, including in Gram-negative bacteria is growing [34]. We also note the high prevalence of fluoroquinolone resistance for some of the syndromes contributing to sepsis mortality, e.g. [35]. Additionally, we found that the share of African Americans in an age group is associated with septicemia mortality rate within that age group for persons aged over 65 years, and 18-49 years in the multivariable analysis, which agrees with the findings that rates of septicemia mortality and hospitalization in African Americans are elevated [29,36]. We also found that the percent of state residents aged 65-74 years living below the poverty level is associated with septicemia mortality rate in that age group.

Our analyses have not found associations between the rates of fluroquinolone prescribing in older adults, as well as the rates of macrolide prescribing and the rates of septicemia mortality. If (like all models) this model is somewhat mis-specified, competition with antibiotics that are more likely to promote sepsis mortality could bias downward the strength of the association between the use of other antibiotics and the rates of sepsis mortality. Additionally, various sources of noise, such as the fact that we related antibiotic prescribing rates in the population overall to rates of septicemia mortality in different age groups, might have reduced the precision of our estimates. We note the high prevalence of resistance to fluoroquinolones for certain syndromes that contribute to sepsis mortality, like urinary tract infections (UTIs) in older adults [31]. Macrolide use could potentially contribute to the rates of sepsis mortality as macrolides are commonly prescribed for the treatment of certain syndromes that are major causes of sepsis, notably respiratory diseases, both chronic [37] and acute, including pneumonia [38]. Overall, the potential benefits of antibiotic stewardship for penicillins in older adults, cephalosporins and fluoroquinolones in younger adults, as well as cephalosporins in persons aged 65-74 years may be the most important findings of this paper, while options for antibiotic replacement require further investigation. We note that recently, US FDA recommended the restriction of fluoroquinolone use for certain conditions (such as uncomplicated UTIs) due to potential adverse effects [39]. Our results suggest that replacement of fluoroquinolones by penicillins and cephalosporins in treating older US adults could potentially lead to increases in septicemia mortality, with that issue needing further study.

Our study has some additional limitations. The antibiotic-sepsis incidence associations we find estimate causal effects only if the model is well-specified and all confounders are accounted for in the analysis. To adjust for potential effects of unmeasured and residual confounding, we included random effects for the ten US Health and Human Services regions, which led to an improvement in the model fits. Further work involving more granular data is needed to ascertain the strength of the associations found in this paper. Data on outpatient antibiotic prescribing in the whole population were used in the regression model for which the outcomes were age-specific rates of septicemia hospitalization. We expect that this source of incompatibility should generally introduce noise into the regression model, reducing precision rather than creating spurious associations.

We believe that despite those limitations, our findings indicate a possible causal association between the use of penicillins and rates of sepsis mortality in older adults, as well as the use of cephalosporins and rates of sepsis mortality in persons aged 18-49 years and 65-74 years, and the use of fluoroquinolones and rates of sepsis mortality in persons aged 18-49 years. We hope that this population-level study would lead to further investigations of the relation between antibiotic prescribing practices and sepsis mortality in different contexts, including the potential effect of antibiotic replacement in the treatment of different syndromes on the rates of sepsis mortality.

## References

[1] Elixhauser A, Friedman B, Stranges E. Septicemia in U.S. Hospitals, 2009: Statistical Brief #122. HealthCare Cost and Utilization Project (HCUP), Agency for Healthcare Research and Quality (AHRQ), 2011. Available from: https://www.hcup-us.ahrq.gov/reports/statbriefs/sb122.pdf

[2] McDermott KW, Elixhauser A, Sun R. Trends in Hospital Inpatient Stays in the United States, 2005–2014. Statistical Brief #225. HealthCare Cost and Utilization Project (HCUP), Agency for Healthcare Research and Quality (AHRQ), 2017. Available from: https://www.hcup-us.ahrq.gov/reports/statbriefs/sb225-Inpatient-US-Stays-Trends.pdf

[3] Iwashyna TJ, Cooke CR, Wunsch H, Kahn JM. Population burden of long-term survivorship after severe sepsis in older Americans. J Am Geriatr Soc. 2012;60(6):1070–7.

[4] Torio CM, Moore BJ. National Inpatient Hospital Costs: The Most Expensive Conditions by Payer, 2013: Statistical Brief #204. HealthCare Cost and Utilization Project (HCUP), Agency for Healthcare Research and Quality (AHRQ), 2016. Available from: https://www.hcup-us.ahrq.gov/reports/statbriefs/sb204-Most-Expensive-Hospital-Conditions.jsp

[5] US CDC. Sepsis. Data & Reports. (2018) Available from: https://www.cdc.gov/sepsis/datareports/index.html

[6] Iwashyna TJ, Cooke CR, Wunsch H, Kahn JM. Population burden of long-term survivorship after severe sepsis in older Americans. J Am Geriatr Soc. 2012;60(6):1070–7.

[7] Rhee C, Murphy MV, Li L, Platt R, Klompas M. Comparison of Trends in Sepsis Incidence and Coding Using Administrative Claims Versus Objective Clinical Data. Clin Infect Dis. 2015;60(1):88–95

[8] Umscheid CA, Betesh J, VanZandbergen C, Hanish A, Tait G, Mikkelsen ME, et al. Development, implementation, and impact of an automated early warning and response system for sepsis. J Hosp Med. 2015;10(1):26–31

[9] Walkey AJ, Lagu T, Lindenauer PK. Trends in sepsis and infection sources in the United States. A population-based study. Ann Am Thorac Soc. 2015;12(2):216–20

[10] Goldstein E, Olesen SW, Karaca Z, Steiner CA, Viboud C, Lipsitch M. Levels of prescribing for four major antibiotic classes and rates of septicemia hospitalization in different US states. bioRxiv 2018. Available from: https://www.biorxiv.org/content/early/2018/08/29/404046

[11] Goldstein E, MacFadden DR, Karaca Z, Steiner CA, Viboud C, Lipsitch M. Antimicrobial resistance prevalence and rates of hospitalization with septicemia in the diagnosis in adults in different US states. bioRxiv 20181. Available from: https://www.biorxiv.org/content/early/2018/08/29/404137

[12] Paul M, Shani V, Muchtar E, Kariv G, Robenshtok E, Leibovici L. Systematic review and meta-analysis of the efficacy of appropriate empiric antibiotic therapy for sepsis. Antimicrob Agents Chemother. 2010;54(11):4851–63

[13] Zilberberg MD, Shorr AF, Micek ST, Vazquez-Guillamet C, Kollef MH. Multi-drug resistance, inappropriate initial antibiotic therapy and mortality in Gram-negative severe sepsis and septic shock: a retrospective cohort study. Crit Care. 2014;18(6):596

[14] Lee YC, Hsiao CY, Hung MC, Hung SC, Wang HP, Huang YJ, et al. Bacteremic Urinary Tract Infection Caused by Multidrug-Resistant Enterobacteriaceae Are Associated With Severe Sepsis at Admission: Implication for Empirical Therapy. Medicine (Baltimore). 2016;95(20):e3694

[15] Lishman H, Costelloe C, Hopkins S, Johnson AP, Hope R, Guy R, et al. Exploring the relationship between primary care antibiotic prescribing for urinary tract infections, Escherichia coli bacteraemia incidence and antibiotic resistance: an ecological study. Int J Antimicrob Agents. 2018 Aug 23. doi:10.1016/j.ijantimicag.2018.08.013

[16] Tacconelli E, De Angelis G, Cataldo MA, Pozzi E, Cauda R. Does antibiotic exposure increase the risk of methicillin-resistant Staphylococcus aureus (MRSA) isolation? A systematic review and meta-analysis. J Antimicrob Chemother. 2008;61(1):26–38

[17] Weber SG, Gold HS, Hooper DC, Karchmer A, Carmeli Y. Fluoroquinolones and the risk for methicillin-resistant Staphylococcus aureus in hospitalized patients. Emerg Infect Dis. 2003;9: 1415 –22

[18] Pouwels KB, Freeman R, Muller-Pebody B, Rooney G, Henderson KL, Robotham JV, et al. Association between use of different antibiotics and trimethoprim resistance: going beyond the obvious crude association. J Antimicrob Chemother. 2018;73(6):1700–1707

[19] Goldstein E, Lee RS, MacFadden DR, Lipsitch M. Outpatient prescribing of four major antibiotic classes and prevalence of antimicrobial resistance in US adults. bioRxiv 2018. Available from: https://www.biorxiv.org/content/early/2018/10/30/456244

[20] López-Mestanza C, Andaluz-Ojeda D, Gómez-López JR, Bermejo-Martín JF. Clinical factors influencing mortality risk in hospital-acquired sepsis. J Hosp Infect. 2018;98(2):194–201

[21] Liu V, Escobar GJ, Greene JD, Soule J, Whippy A, Angus DC, et al. Hospital deaths in patients with sepsis from 2 independent cohorts. JAMA. 2014;312(1):90–2.

[22] US CDC Wonder. Multiple Cause of Death, 1999-2016 Request. Available from: https://wonder.cdc.gov/mcd-icd10.html

[23] Epstein L, Dantes R, Magill S, Fiore A. Varying Estimates of Sepsis Mortality Using Death Certificates and Administrative Codes—United States, 1999–2014. MMWR Morb Mortal Wkly Rep 2016;65:342–345.

[24] US CDC. Antibiotic Resistance Patient Safety Atlas. Outpatient Antibiotic Prescription Data. Available from: https://gis.cdc.gov/grasp/PSA/indexAU.html

[25] United States Census Bureau. MedIAN HOUSEHOLD INCOME (IN 2011 INFLATION-ADJUSTED DOLLARS) - United States - States; and Puerto Rico: Households. 2011 American Community Survey 1-Year Estimates (2011). Available from: https://factfinder.census.gov/faces/tableservices/jsf/pages/productview.xhtml?src=bkmk

[26] National Oceanic and Atmospheric Administration (NOAA). National Climatic Data. Available from: https://www7.ncdc.noaa.gov/CDO/CDODivisionalSelect.jsp#

[27] US Centers for Disease Control and Prevention. Bridged-Race Population Estimates, 1990-2016 data request. Available from: https://wonder.cdc.gov/Bridged-Race-v2016.HTML

[28] US Census Bureau. Current Population Survey (CPS) (2018). CPS Table Creator. Available from: https://www.census.gov/cps/data/cpstablecreator.html

[29] Wang HE, Devereaux RS, Yealy DM, Safford MM, Howard G. National variation in United States sepsis mortality: a descriptive study. Int J Health Geogr. 2010;9:9

[30] MacFadden DR, McGough SF, Fisman D, Santillana M, Brownstein JS. Antibiotic resistance increases with local temperature. Nature Climate Change. 2018;8: 510–514

[31] Morrill HJ, Morton JB, Caffrey AR, Jiang L, Dosa D, Mermel LA, et al. Antimicrobial Resistance of Escherichia coli Urinary Isolates in the Veterans Affairs Health Care System. Antimicrob Agents Chemother. 2017;61(5) pii: e02236-16

[32] Cheng MP, René P, Cheng AP, Lee TC. Back to the Future: Penicillin-Susceptible Staphylococcus aureus. Am J Med. 2016;129(12):1331–1333

[33] Baquero, F., and J. Blazquez. Evolution of antibiotic resistance. Trends in Ecology and Evolution 1997;12:482-487

[34] Park, SH. Third-generation cephalosporin resistance in gram-negative bacteria in the community: a growing public health concern. Korean J Intern Med. 2014;29(1):27– 30

[35] Bidell MR, Palchak M, Mohr J, Lodise TP. Fluoroquinolone and Third-Generation-Cephalosporin Resistance among Hospitalized Patients with Urinary Tract Infections Due to Escherichia coli: Do Rates Vary by Hospital Characteristics and Geographic Region? Antimicrob Agents Chemother. 2016;60(5):3170–3

[36] Barnato AE, Alexander SL, Linde-Zwirble WT, Angus DC. Racial variation in the incidence, care, and outcomes or severe sepsis: analysis of population, patient, and hospital characteristics. Am J Respir Crit Care Med. 2008;177:279–284

[37] Suresh Babu K, Kastelik J, Morjaria JB. Role of long term antibiotics in chronic respiratory diseases. Respir Med. 2013;107(6):800–15.

[38] Mandell LA, Wunderink RG, Anzueto A, Bartlett JG, Campbell GD, Dean NC, et al. Infectious Diseases Society of America/American Thoracic Society consensus guidelines on the management of community-acquired pneumonia in adults. Clin Infect Dis. 2007;44 Suppl 2:S27–72

[39] US Food and Drug Administration. FDA updates warnings for fluoroquinolone antibiotics. July 2016. Available from: https://www.fda.gov/NewsEvents/Newsroom/PressAnnouncements/ucm513183.ht m

